# CD99Targeted Irinotecan Containing Nanoparticles Show Twenty-Fold Greater Anti-Tumor Effect Than Free Irinotecan For Treatment of Ewing Sarcoma

**DOI:** 10.64898/2026.01.06.697855

**Authors:** Hyung-Gyoo Kang, Sheetal Mitra, Bryon Upton, Jean Parmentier, Anahit Hovsepyran, Jon Nagy, Timothy J. Triche

## Abstract

**Purpose:** To assess the antitumor efficacy, pharmacokinetics, and safety of NV103, a CD99-targeted liposomal irinotecan nanoparticle, in a preclinical Ewing sarcoma model.

**Experimental Design:** NV103 is a CD99-antibody targeted irinotecan containing nanoparticle, engineered to selectively deliver irinotecan to CD99 expressing tumor cells. *In vitro* studies measured binding, internalization, and cytotoxic IC_50_ values in several Ewing sarcoma cell lines. *In vivo*, mice bearing xenografts derived from treatment-naïve and chemoresistant Ewing lines were treated. NV103 was compared with free irinotecan, untargeted nanoparticles, and Onivyde^TM^ at multiple dosages. Plasma pharmacokinetics of irinotecan and SN-38, biodistribution of the nanoparticles, and toxicity (body weight, organ function, and hematology) were assessed.

**Results:** NV103 bound selectively to tumor cells (>80× over control), was rapidly internalized, and showed enhanced potency *in vitro* (IC_50_ ≈ 3–4 nM at 0.5–1 h). *In vivo*, biweekly dosing at 5 mg/kg resulted in full tumor regression sustained for 140 days, even after stopping treatment at day 70. Effective suppression and survival benefit were observed at doses as low as 1 mg/kg; the ED_50_ was estimated to be between 1–2.5 mg/kg versus 50mg/kg for free irinotecan. In a chemoresistant Ewing tumor cell line, NV103 induced similar tumor-free remission. Pharmacokinetics revealed prolonged and elevated plasma levels of irinotecan with NV103 versus free drug. No systemic toxicity was detected at doses of 10 mg/kg. Biodistribution showed tumor-preferential accumulation.

**Conclusions:** NV103 displays potent and durable antitumor activity in Ewing sarcoma at low doses with no toxicity and favorable pharmacokinetics. These findings support further development for clinical translation.

**Translational Relevance:** Although Irinotecan has shown activity against Ewing sarcoma, its clinical utility is limited by systemic toxicity and poor tumor selectivity. NV103, a CD99-targeted nanoparticle formulation of irinotecan delivers irinotecan selectively to Ewing sarcoma cells. CD99 is a surface antigen that is highly expressed in Ewing sarcoma cells but largely absent from irinotecan-sensitive organs like liver, kidney, and bone marrow. In preclinical xenograft models, NV103 induced complete and durable tumor regression at doses >10 fold lower than those of free irinotecan or untargeted nanoparticles, with no detectable systemic toxicity. These findings suggest that NV103 is a promising translational therapeutic agent that enhances the therapeutic index of irinotecan and other cytotoxic agents. This platform offers a broadly adaptable approach to tumor-specific drug delivery, which could significantly improve treatment outcomes and reduce long-term toxicity in children and young adults with Ewing sarcoma and other solid tumors.

## Introduction

Ewing sarcoma is a malignant bone and soft tissue tumor that primarily affects adolescents and young adults. It is distinct among sarcomas that lack a known normal tissue counterpart, no animal counterpart, or an intrinsic metastatic phenotype (Supplemental Figure 1). As first noted by James Ewing in 1921, local therapies alone are invariably ineffective and systemic approaches are required for meaningful disease control (1, 2). The introduction of multi-agent chemotherapy in the 1960s, particularly combinations such as vincristine, doxorubicin, cyclophosphamide, ifosfamide, and etoposide (VAC/IE), marked the first major therapeutic breakthrough (3–5). However, despite decades of clinical trials, survival rates have remained static at approximately 70% for patients with localized disease and less than 30% for those with metastases at diagnosis (6, 7). Recent efforts to incorporate molecularly targeted therapies, such as IGF-1R inhibitors, tyrosine kinase inhibitors (e.g., regorafenib and cabozantinib), EWS-FLI1 fusion inhibitors (e.g., TK216), and CAR-T cell therapy, have not produced significant improvements in overall survival (8, 9). Therefore, there is a critical unmet need for novel therapies that are both more effective and less toxic.

Nanoparticle-based drug delivery platforms, particularly liposomal formulations, have demonstrated clinical success in other tumors by improving the therapeutic index of chemotherapeutic agents (10–12). However, most of these systems rely on passive accumulation and lack tumor-specific targeting, leading to suboptimal efficacy and off-target toxicities (13–15). Achieving true tumor selectivity requires the active targeting of antigens preferentially or uniquely expressed on tumor cells (16, 17).

Ewing sarcoma has demonstrated pronounced sensitivity to irinotecan, a topoisomerase I inhibitor, in preclinical studies and salvage regimens for relapsed or metastatic disease (18). While irinotecan encapsulated in untargeted liposomal nanoparticles improves delivery and reduces systemic exposure, these formulations have not been proven curative (19). We hypothesized that the selective delivery of irinotecan using actively targeted nanoparticles could significantly enhance tumor cell killing while reducing toxicity to normal tissues.

The challenge in implementing targeted delivery lies in identifying an appropriate tumor-associated surface antigen (20, 21). Similar to HER2-targeted therapy in breast cancer (e.g., trastuzumab) (22, 23), Ewing sarcoma has long been known to express high levels of CD99, also known as p30/32MIC2 (24). CD99 is a surface glycoprotein strongly expressed in nearly all Ewing sarcoma cells and only minimally expressed in normal tissues, particularly the liver, kidney, intestine, and hematopoietic progenitors—tissues commonly affected by irinotecan toxicity (25, 26). This differential expression pattern suggests that CD99 is an ideal candidate for targeted therapeutic delivery (27, 28).

In this study, we developed and tested a novel CD99-targeted irinotecan-loaded nanoparticle (designated NV103) (29). We evaluated its efficacy in vitro and in vivo using immunodeficient mice xenografted with multiple well-characterized Ewing sarcoma cell lines, with a focus on antitumor efficacy, tissue specificity, pharmacokinetics, and toxicity. Our findings indicate that CD99-targeted nanoparticles may offer a promising therapeutic strategy for overcoming the limitations of conventional irinotecan therapy in Ewing sarcoma.

## Results

### Nanoparticle Characterization, Targeting Efficiency, Cellular Uptake & Trafficking

Hybrid polymerized lipid nanoparticles (HPLNs) were formulated using a mixture of hydrogenated soy PC (where the primary component is distearoylphosphatidylcholine (DSPC)), cholesterol, polyethylene glycol-distearoylphosphatidyl ethanolamine (m-PEG2000-DSPE), N-(methoxy(polyethylene glycol)-2000)-10-12-pentacosadiynamide (m-Peg_2000_-PCDA), and N-(5’-hydroxy-3’-oxypentyl)-10-12-pentacosadiynamide (h-Peg_1_-PCDA), as illustrated in Supplement Figure 2A. The inclusion of PCDA enabled UV cross-linking, which conferred structural stability and enhanced the drug metabolism and pharmacokinetic (DMET) properties. This formulation also supported the active loading of irinotecan, resulting in substantially higher drug concentrations compared to the passive loading methods (13). The cross-linked PCDA component also imparted intrinsic red fluorescence, enabling visualization without additional labeling.

Antibody conjugation was achieved by coupling the reduced antibodies to maleimide-terminated DSPE micelles via sulfhydryl-maleimide chemistry. These antibody-bearing micelles were then fused with irinotecan-loaded HPLNs to generate targeted nanoparticles. The resulting antibody-conjugated nanoparticles were approximately 90 nm in diameter, as shown in Supplementary Figure 2B, and featured a hollow aqueous core (Supplementary Figure 2C). Time-course nanoparticle tracking of NV103 demonstrated that nanoparticle size remained unchanged even after 180 days (Supplementary Figure 2D).

Immunoblotting demonstrated that all six Ewing’s tumor cell lines express CD99 at high levels (Figure 1A). To evaluate the targeting specificity, antibody-conjugated nanoparticles were incubated with six Ewing sarcoma cell lines. All six cell lines demonstrated robust binding of the antibody-labeled nanoparticles, exhibiting >80-fold greater fluorescence intensity than unlabeled controls, which showed minimal binding (Figures 1B). Within 4 h of incubation, intense red fluorescence was observed on both the cell surface and intracellular compartments, consistent with the rapid cellular uptake and internalization of the nanoparticles (Figure 1C). The antibody density on the nanoparticle surface was modulated by varying the ratio of the antibody micelles to HPLNs. Binding studies indicated optimal cellular interaction at an antibody concentration of ∼28 µg/mg of HPLN, corresponding to an estimated eight antibody molecules per nanoparticle (Figure 1D). As shown in Supplemental Figure 3, NV103 treatment induced apoptosis by activating the caspase-dependent cell death pathways.

**Figure 1.**
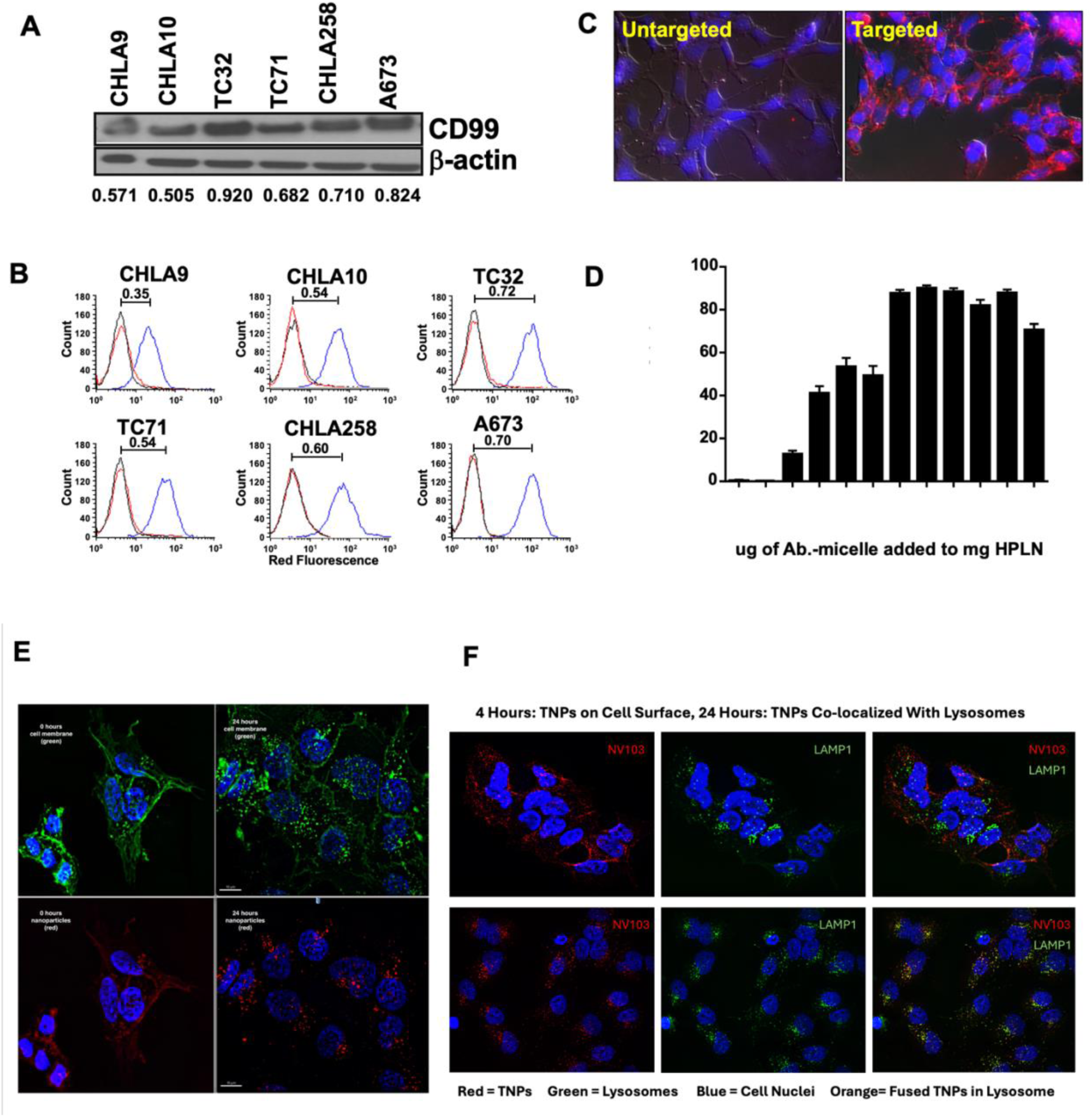
Binding and Internalization of Antibody-Labelled Nanoparticles in Ewing Sarcoma Cell Lines. (A) CD99 expression in EFT cell lines accessed by immunoblotting. Relative band intensity was obtained by normalization with the expression of b-actin. (B) Flow cytometry analysis of nanoparticle binding. Antibody-labelled nanoparticles (blue) and unlabeled nanoparticles (red). (C) Fluorescence microscopy images captured at 4 hours post-incubation of nanoparticles. (D) Optimization of Antibody Density on Hybrid Polymerized Lipid Nanoparticles (HPLNs) for Enhanced Cell Binding. Antibody density on HPLNs was optimized by varying the amounts of antibody-conjugated micelles fused with the nanoparticles. Cell binding of HPLNs was detected with Flow cytometry. (E) Time course of nanoparticle binding, internalization, and drug release in Ewing tumor cells. Cellular membranes (green) and nanoparticles (red) were visualized at 0 and 24 hours. (F) To assess intracellular trafficking, lysosomes were stained green and compared to nanoparticle localization at 4 and 24 hours. To visualize nuclear morphological changes induced by irinotecan, DAPI staining was performed concurrently.

Time-course analysis was conducted to assess nanoparticle binding, internalization, and intracellular drug release. In the initial imaging at 0 h, red fluorescence from the nanoparticles co-localized predominantly with green-stained cell membranes, indicating surface binding (Figure 1E). After 24 h, both the membrane stain and nanoparticle signal exhibited a punctate, intracellular distribution consistent with endophagocytic internalization, as opposed to direct cell membrane fusion and diffuse cytoplasmic distribution.

To investigate the subcellular fate of the internalized nanoparticles, lysosomes were stained green and imaged 4- and 24-hours post-incubation (Figure 1F). At 4 h, minimal overlap was observed between the nanoparticle and lysosomal signals, suggesting limited lysosomal localization. However, after 24 h, significant colocalization was evident, indicating that nanoparticles had localized to lysosomal compartments via endophagocytosis, as opposed to direct membrane fusion, as observed with unpolymerized liposomal nanoparticles.

Evidence of intracellular drug release was supported by nuclear morphology changes detected by DAPI staining. At 24 h, the nuclei exhibited heterogeneous chromatin condensation compared with the homogenous DAPI staining observed at 4 h. These nuclear alterations were consistent with irinotecan-induced DNA damage, suggesting the successful intracellular release of the drug payload from the nanoparticles. As shown in Supplemental Figure 4, in NV103-treated cells, the appearance of the apoptotic markers Annexin V (early marker) and caspase 3 (late marker) precedes the nuclear apoptotic changes seen on cellular imaging.

### Comparative IC_50_ of Irinotecan Formulations

To evaluate the relative potency of free irinotecan, liposomal irinotecan (Onivyde^TM^), untargeted HPLN-irinotecan (HPLN-Ir), and CD99-targeted HPLN-irinotecan (NV103), *in vitro* cytotoxicity assays were performed across three exposure durations: 0.5 hours, 1 hour, and 72 hours. At both 0.5 and 1 h exposure times, NV103 demonstrated the greatest potency, with IC_50_ values of 8.30 μM and 0.795 μM, respectively (Figure 2). In contrast, the other formulations required higher concentrations to achieve comparable cell death at these early time points.

**Figure 2.**
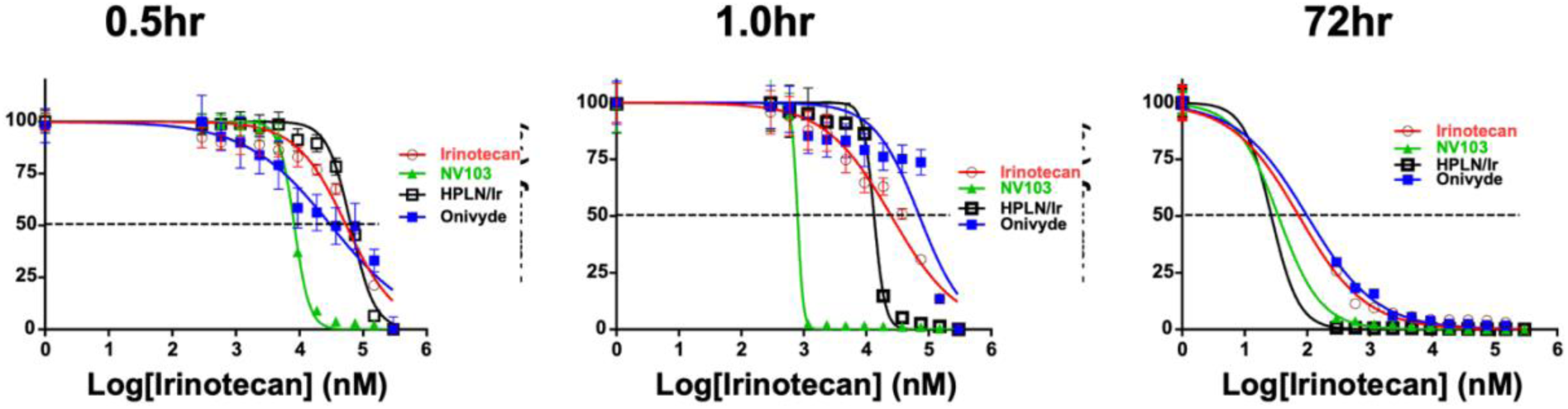
Determination of relative IC_50_ values for irinotecan in different formulations using an *in vitro* cell viability assay. To assess the cytotoxic efficacy of free irinotecan, Onivyde^TM^, untargeted HPLN-Ir, and Cd99-targeted HPLNs (NV103), Ewing sarcoma cells were exposed to each formulation for 0.5 hours, 1 hour, and 72 hours.

Following 72 hours of continuous exposure, both NV103 and untargeted HPLN-Ir achieved an IC_50_ of 34.58 and 26.74nM, respectively indicating that prolonged incubation allowed equilibration and effective delivery of irinotecan irrespective of targeting. However, at early time points, the superior cytotoxicity of NV103 was consistent with enhanced nanoparticle-cell interaction and rapid uptake, as previously demonstrated in binding assays (Figure 1C).

These findings support the conclusion that CD99-targeted nanoparticles mediate a more efficient early intracellular delivery of irinotecan, resulting in lower IC_50_ values under limited exposure conditions. This suggests a potentially lower effective dose (ED_50_) may be achievable in *in vivo* models using NV103.

### In Vivo Efficacy and Dose Escalation and Comparative Efficacy Studies of NV103

The therapeutic efficacy of CD99-targeted irinotecan-loaded nanoparticles (NV103) was compared to that of untreated controls in Ewing sarcoma xenograft-bearing mice. The animals received a biweekly intravenous dose of 5 mg/kg (drug-equivalent) of irinotecan. As shown in Figure 3A, the group treated with NV103 exhibited complete suppression of tumor growth.

**Figure 3.**
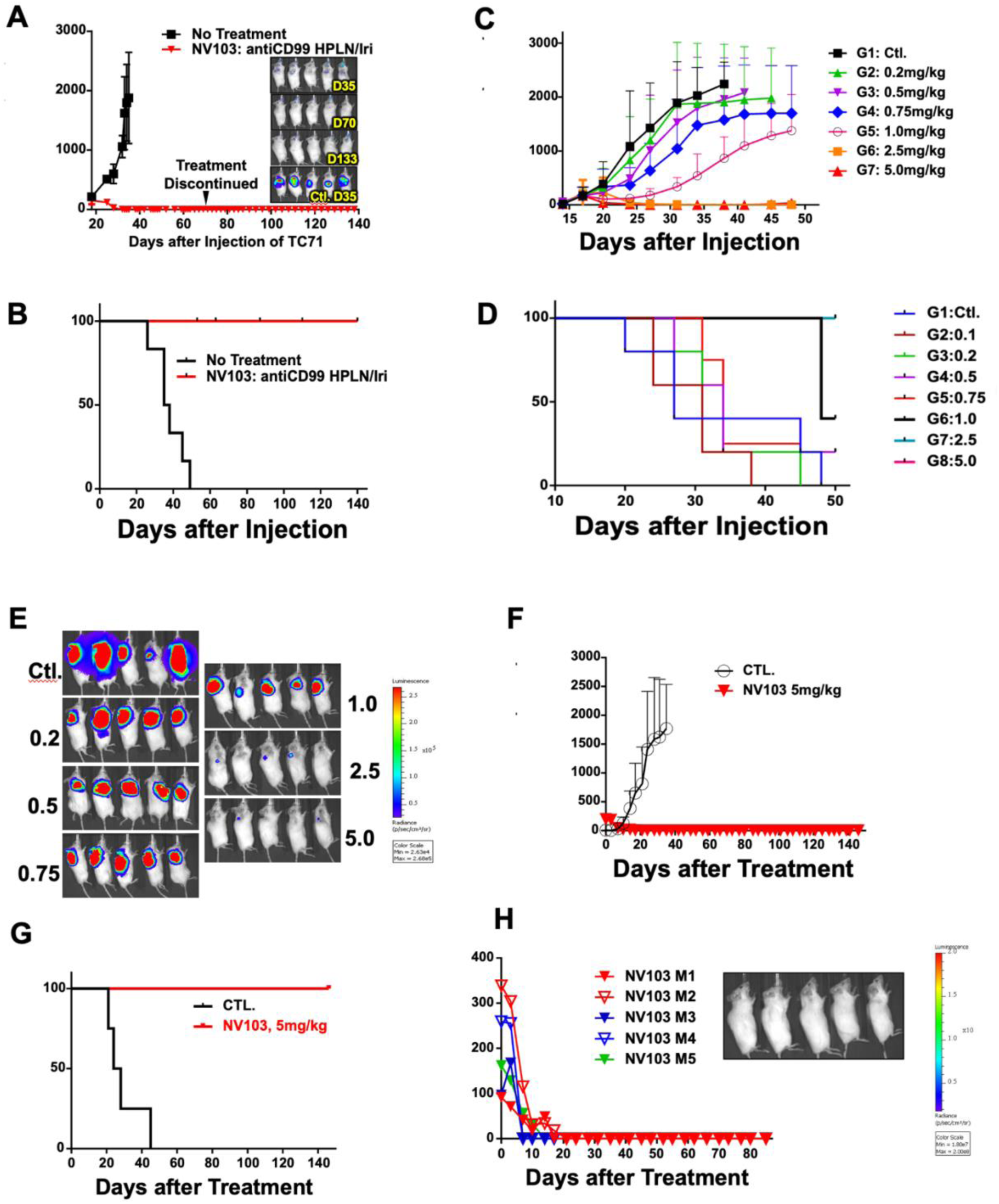
In vivo efficacy of CD99-targeted irinotecan-loaded nanoparticles (NV103) in Ewing sarcoma xenograft model. (A) Mice bearing Ewing tumor xenografts were treated twice weekly with 5 mg/kg of irinotecan-loaded nanoparticles Irinotecan treatment was terminated at day 72. (B) Kaplan-Meier survival analysis of each treatment cohort (C) Dose escalation studies to determine the ED50 of targeted irinotecan-containing nanoparticles and comparison with untargeted formulations. To identify the ED50 of CD99-targeted irinotecan-loaded nanoparticles, mice bearing Ewing tumor xenografts were treated twice weekly with doses ranging from 0.2 to 5 mg/kg. Tumor size was measured by caliper measurement. (D) Kaplan-Meier survival analysis of each treatment cohort (E) Bioluminescent imaging of luciferase-expressing Ewing tumors treated with different dose of irinotecan-loaded nanoparticles on Day20. (F) Efficacy of CD99-targeted irinotecan-loaded nanoparticles (NV103) in overcoming tumor growth in a treatment-resistant Ewing sarcoma (CHLA10) xenograft model. Mice bearing CHLA10 Ewing sarcoma xenografts were treated with 5 mg/kg NV103 twice weekly. Tumor volume was measured by calipers. (G) Kaplan-Meier survival analysis of treated and control cohorts. (H) Bioluminescent imaging of luciferase-expressing CHLA10 treated with 5 mg/kg NV103 twice weekly on Day 55.

Notably, mice treated with NV103 at 5 mg/kg exhibited no tumor growth over the entire 140-day period following xenograft, even after treatment was stopped on day 70. This result was further supported by Kaplan–Meier survival analysis (Figure 3B), where all animals in the control group reached IACUC-defined humane endpoints due to excessive tumor burden, while all animals in the NV103 group remained alive and tumor-free through day 140.

Bioluminescence imaging of luciferase-expressing Ewing sarcoma tumors confirmed these findings. By day 35, minimal signal was detectable in NV103-treated animals, and no bioluminescence was observed at subsequent time points (Figure 3A), indicating complete tumor eradication. These data demonstrated the superior *in vivo* efficacy of CD99-targeted irinotecan nanoparticles (NV103).

To determine the estimated effective dose (ED_50_) of CD99-targeted irinotecan-loaded nanoparticles (NV103), a dose escalation study was performed. Mice bearing Ewing sarcoma xenografts were treated biweekly with doses ranging from 0.2 to 5 mg/kg (n = 5 per group). As shown in Figure 3C and 3E, tumor suppression was dose-dependent, with increasing inhibition observed at 0.2, 0.5, 0.75, and 1.0 mg/kg. Complete tumor suppression was achieved at doses of 2.5 mg/kg and 5 mg/kg, and all mice in those two groups were still alive at day 50 (Figure 3D). Based on these data, the ED_50_ of NV103 was estimated to lie between 1.0 and 2.5 mg/kg.

To evaluate whether CD99-targeted irinotecan nanoparticles (NV103) could overcome resistance in relapsed Ewing sarcoma, we used CHLA-10, a cell line derived from a patient whose tumor progressed despite standard 5-agent VAC/IE chemotherapy. Five NSG mice bearing CHLA-10 xenografts were treated with NV103 at 5 mg/kg twice weekly. As shown in Figure 3F, all NV103-treated mice exhibited complete and durable tumor regression within 20 days of treatment initiation. No residual tumor signal was detected on bioluminescence imaging at this time point (Figure 3G). In contrast, all untreated control mice demonstrated progressive tumor growth, requiring sacrifice by day 42. All NV103-treated mice remained tumor-free and survived the 150-day observation period, with no evidence of recurrence (Figure 3H).

### Dose Escalation and Intensification of NV103

To compare efficacy with existing therapies, a second dose-escalation experiment was conducted, evaluating Onivyde^TM^ (a clinically available untargeted liposomal irinotecan), untargeted HPLN-irinotecan, and CD99-targeted HPLNs, each administered at 1, or 10 mg/kg. At the 1 mg/kg dose level, only the HPLN-based formulations conferred a survival benefit: untargeted HPLNs achieved 100% survival and CD99-targeted HPLNs achieved 80% survival at day 36 post-treatment (Figure 4A). One out of 5 died of an undetermined cause, not from the tumor. At the time of death, a minimal amount of tumor was detected. In contrast, all mice treated with Onivyde^TM^ showed progressive tumor growth, with no survival beyond day 36, similar to the untreated controls (median survival: 34 days).

**Figure 4.**
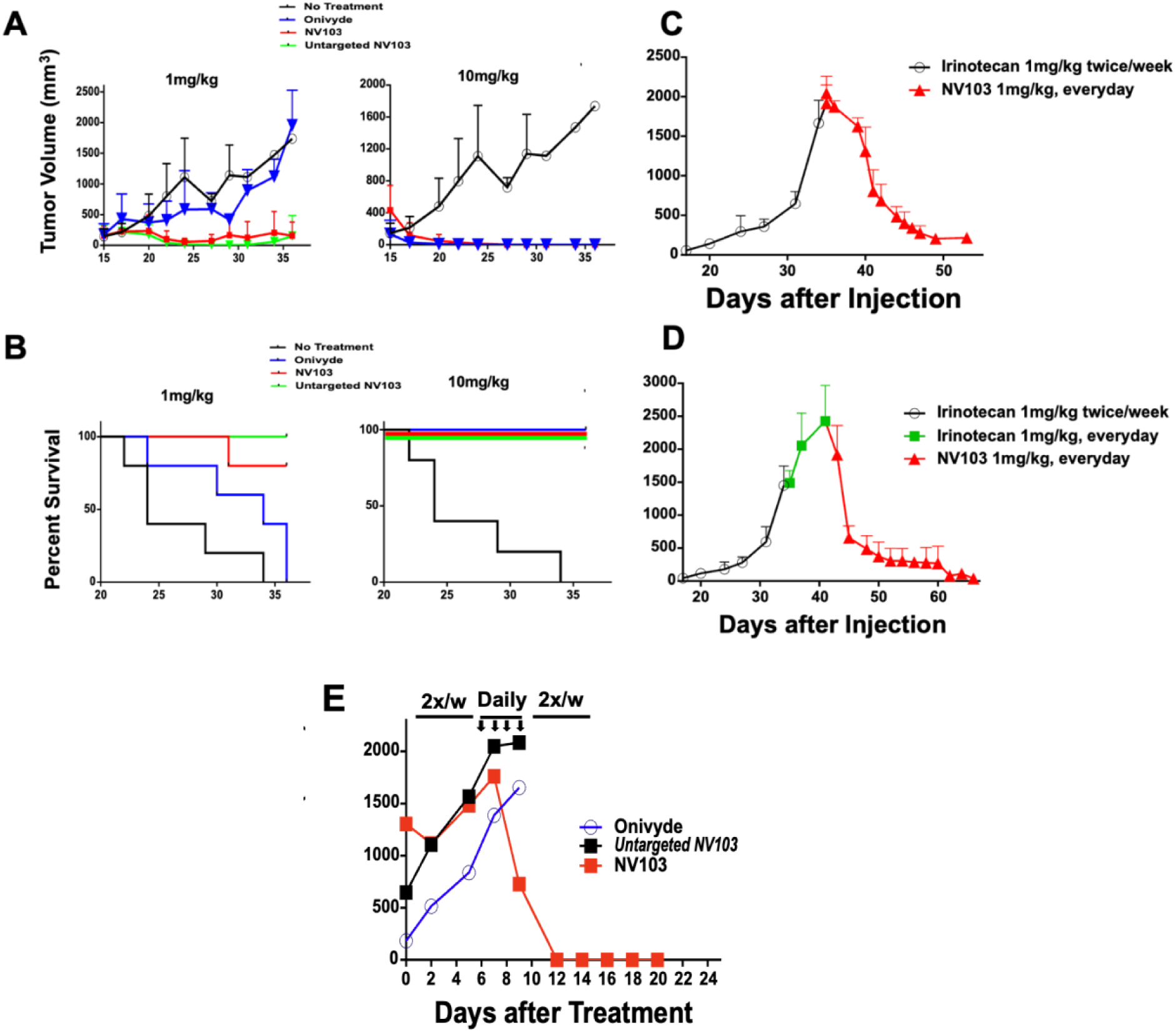
Dose escalation and intensification studies of targeted irinotecan-containing nanoparticles and comparison with untargeted formulations. (A) Dose escalation experiment comparing Onivyde^TM^, untargeted irinotecan-loaded HPLNs, CD99-targeted HPLNs (NV103), and untreated control animals. (B) Kaplan-Meier survival analysis of each treatment cohort (C) To assess whether dose intensification at a lower dose (1 mg/kg) could enhance tumor suppression, mice were initially treated with 1 mg/kg of free irinotecan twice weekly until tumor volumes exceeded 1,500 mm³. At this point, mice were transitioned to daily 1 mg/kg dosing of free irinotecan. After 6 days of irinotecan treatment, the mice were switched to a daily schedule of NV103 (1mg/kg). (D) Mice were initially treated with 1 mg/kg of free irinotecan twice weekly until tumor volumes exceeded 1,500 mm³. At this point, mice were switched to daily 1 mg/kg dosing of NV103. (E) Efficacy comparison of Onivyde^TM^, untargeted irinotecan-loaded HPLNs, and NV103 in the same model. Animals were treated twice weekly with 1 mg/kg until tumor volumes exceeded 1,500 mm³, followed by daily 1 mg/kg dosing for four days, after which treatment reverted to twice-weekly dosing.

At a dose of 1 mg/kg, all HPLN-treated animals, both targeted and untargeted, showed significant tumor suppression and over 80% survival by day 36. These findings indicate that the HPLN formulations exhibit significantly greater in vivo potency than Onivyde^TM^, with at least a five-fold improvement in efficacy.

To evaluate whether dose intensification at a subtherapeutic level (1 mg/kg) could improve tumor suppression, mice bearing Ewing sarcoma xenografts were initially treated with 1 mg/kg of free irinotecan administered twice weekly. Once tumors exceeded 1,500 mm³, animals were transitioned to daily dosing with either free irinotecan or CD99-targeted nanoparticles (NV103) at a dose of 1 mg/kg. Mice receiving daily NV103 demonstrated rapid and pronounced tumor regression (Figure 4C), whereas those receiving free irinotecan showed no reduction in tumor burden (Figure 4D).

Subsequently, free irinotecan-treated mice were switched to daily dosing with NV103 at 1 mg/kg. Significant tumor regression was observed, and by 10 days, tumor volumes had decreased to less than 5% of their original size. By day 70, little or no detectable tumors remained in any NV103-treated animals (Figure 4D [panel reference]).

A parallel experiment was conducted to compare the effects of intensified dosing of different irinotecan formulations. Individual mice were treated with Onivyde^TM^, untargeted HPLN-Ir, or NV103 (1 mg/kg) twice weekly. Once tumors exceeded 1,500 mm³, daily dosing was initiated for four consecutive days, followed by a return to the original twice-weekly schedule. Only mice treated with NV103 showed complete tumor regression. Neither Onivyde^TM^ nor untargeted HPLNs resulted in meaningful tumor suppression, even with dose intensification (Figure 4E).

### Plasma Pharmacokinetics and Metabolism

To assess the systemic availability of NV103 compared to free irinotecan, we performed a pharmacokinetic time-course analysis measuring the plasma concentrations of both irinotecan and its active metabolite, SN-38, in normal mice. As shown in Figure 5, NV103 exhibited markedly superior pharmacokinetic properties compared to those of free irinotecan.

**Figure 5.**
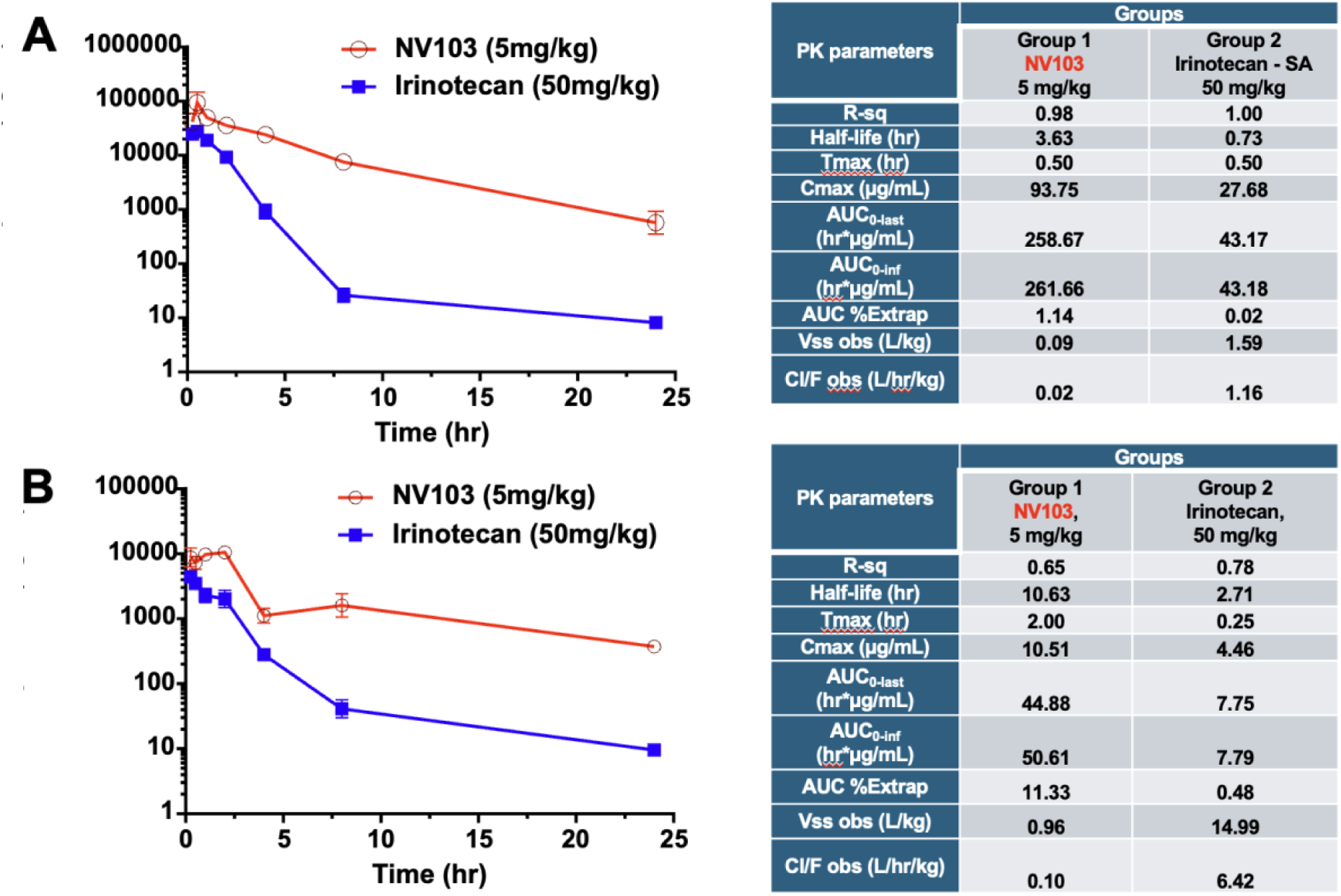
Plasma Pharmacokinetics and Metabolism of NV103-Delivered Irinotecan and SN-38. Time-course analysis of plasma concentrations of irinotecan (CPT-11) (A) and its active metabolite SN-38 (B) was performed in mice following administration of a single 5 mg/kg dose of NV103 nanoparticles or a 50 mg/kg dose of free irinotecan

Despite being administered at one-tenth of the dose (5 mg/kg for NV103 vs. 50 mg/kg for free irinotecan), NV103 achieved equivalent initial plasma irinotecan concentrations (∼100 µg/mL). Notably, NV103 maintained significantly higher plasma levels over time, with concentrations remaining above 1 µg/mL at 25 h post-injection. In contrast, free irinotecan levels declined rapidly, falling below 8 ng/mL at the same time point, with a greater than 100-fold difference.

Pharmacokinetic analysis (Figure 5, tables) confirmed this sustained exposure, with NV103 demonstrating a six-fold greater area under the curve (AUC) and a five-fold longer half-life than free irinotecan. These findings suggest that nanoparticle encapsulation dramatically enhances drug retention and systemic exposure.

The plasma levels of the active metabolite SN-38 followed a similar trend. While the initial SN-38 concentrations were approximately ten-fold lower than irinotecan for both formulations, NV103 sustained higher levels over time. The half-life of SN-38 was three times longer in NV103-treated mice, and its AUC was significantly greater, despite the lower starting dose. These pharmacokinetic advantages are consistent with the enhanced in vitro cytotoxicity and in vivo tumor regression observed with NV103, supporting its superior therapeutic profile compared with free irinotecan (Supplemental Table 1-4).

### Toxicity, Safety and Tissue Distribution of NV103

To evaluate the safety of CD99-targeted irinotecan nanoparticles (NV103), dose-related toxicity was assessed in multiple *in vivo* studies. No signs of treatment-related toxicity were observed in any cohort treated with doses of up to 10 mg/kg administered twice weekly. As shown in Figure 6A, tumor-bearing NSG mice maintained stable body weight during six weeks of bi-weekly treatment at 10 mg/kg. Similarly, C57BL/6 mice showed no abnormalities in liver or kidney function or hematologic parameters following twice-weekly NV103 administration at 10 mg/kg for two weeks.

**Figure 6.**
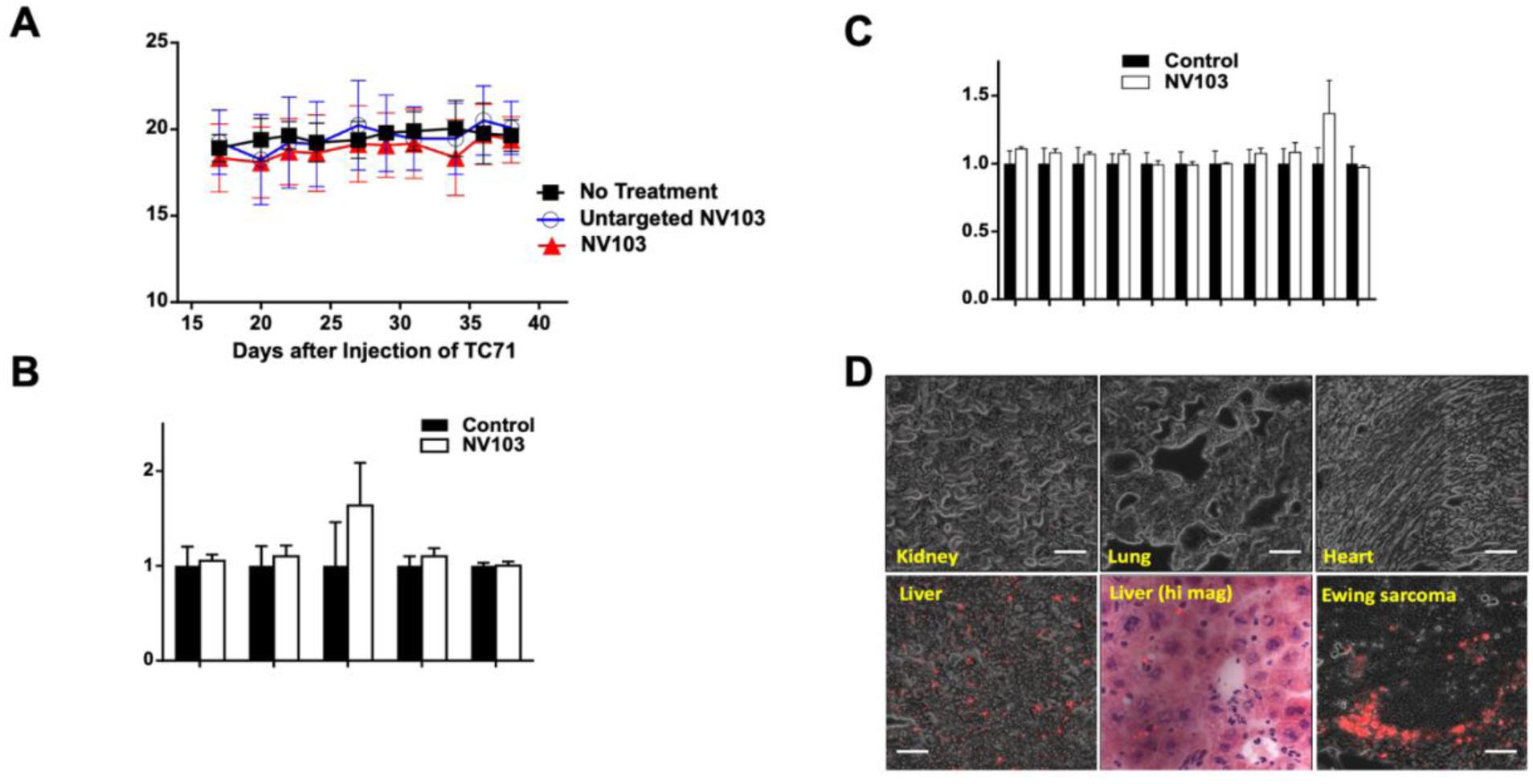
Toxicity evaluation and tissue distribution of NV103. Normal C57BL Mice were dosed twice weekly with 10 mg/kg NV103 for six weeks. (A) Body weight (B) Liver or kidney function (C) Hematopoiesis in treated animals. (D) Distribution of NV103 in tissues and tumor. Fluorescence microscopy images of formalin-fixed, paraffin-embedded tissue sections reveal the biodistribution of NV103. Red fluorescent dots indicate areas of nanoparticle accumulation.

To characterize the biodistribution of CD99-targeted NV103 nanoparticles, fluorescence microscopy was performed on formalin-fixed paraffin-embedded tissues harvested from treated animals. The tissues examined included known sites of nanoparticle sequestration (liver and kidney), tissues with low expected uptake (lung, myocardium), and tumors. Image analysis was used to quantify nanoparticle accumulation, which was visualized as red fluorescent signals.

As shown in Figure 6D, high levels of NV103 accumulation were observed in the tumor tissue, where nanoparticles were present as numerous punctate red fluorescent signals. In contrast, the kidney, lung, and heart tissues exhibited negligible nanoparticle presence, with minimal or no visible fluorescence. In the liver, nanoparticle accumulation was predominantly localized in the portal regions and hepatic sinusoids, with little to no uptake observed in hepatocytes. This pattern is consistent with uptake by Kupffer cells and binding to CD99-expressing lymphocytes in the portal tracts.

Despite liver accumulation, there was no histological or functional evidence of hepatic or hematological toxicity. Hematopoiesis remained intact, and differential blood counts were stable throughout the treatment, as previously shown in Figure 6 B and C, supporting the favorable safety profile of NV103.

## Discussion

The findings of this study provide compelling evidence that Ewing sarcoma can be effectively treated with CD99-targeted irinotecan-loaded nanoparticles (NV103). Complete and sustained tumor regression was achieved in xenograft models with NV103 at doses as low as 1 mg/kg and consistently at 2.5 mg/kg. This efficacy was not observed with commercially available untargeted formulations, such as Onivyde^TM^, nor with untargeted high-payload lipid nanoparticles (HPLN/irinotecan) at the same dose, highlighting the critical contribution of CD99-mediated tumor targeting (19).

The data support the conclusion that the minimum effective dose (MED) of NV103 is approximately 2 mg/kg, nearly 50-fold lower than that required for free irinotecan, and at least five-fold lower than that of untargeted nanoparticle formulations. These findings underscore the importance of antibody-directed targeting for improving therapeutic specificity and potency. Targeting CD99, a surface antigen that is highly expressed on Ewing sarcoma cells, facilitates tumor localization and enhances the intracellular delivery of drug payloads, contributing to the superior antitumor activity of NV103.

Although improved tumor targeting is central to NV103 efficacy, the therapeutic advantage likely arises from multiple synergistic mechanisms. These include prolonged plasma persistence, elevated systemic drug concentrations, enhanced tumor retention, and intracellular delivery via endocytosis (27, 29, 30). Notably, targeted delivery also circumvents common resistance pathways such as MDR1-mediated drug efflux. NV103-mediated delivery resulted in detectable DNA damage within eight hours of administration, consistent with the early and effective cytotoxic action. Additionally, NV103 demonstrated exceptional efficacy against the chemotherapy-resistant Ewing sarcoma model, CHLA-10, which is known for its resistance to conventional treatments and is commonly used to evaluate new therapies. CHLA-10 is characterized by TP53 mutation, polyploidy with pseudo-tetrasomy, and multiple chromosomal duplications, and displays markedly higher proliferation rates in vitro compared to the treatment-naïve CHLA-9 line from the same patient, which is diploid and TP53 wild-type (31, 32).

Collectively, these properties contribute to the enhanced therapeutic index of NV103, achieving robust tumor eradication with minimal systemic toxicity.

Notably, NV103 demonstrated an excellent safety profile. Across all dose levels tested (0.2–10 mg/kg), there was no observable treatment-related toxicity, even at the upper dose range where the efficacy plateaued. Parallel studies in normal C57BL/6 and NSG mice revealed no adverse effects on body weight, organ function (liver and kidney), or hematopoiesis. In contrast, free irinotecan is known to induce dose-limiting toxicities at clinically relevant doses (mouse equivalent ∼50 mg/kg) and is ineffective in achieving tumor ablation in these models. This highlights the limitations of conventional irinotecan administration and reinforces the clinical relevance of NV103’s improved safety and efficacy profiles. It is not feasible to compare free irinotecan to nanoparticle-encapsulated irinotecan to assess toxicity for several reasons, including pharmacodynamics, including distribution, metabolism, excretion, and specific tissue toxicity (DMET), which are completely different for free drug compared to nanoparticle-encapsulated drug. (33). All nanoparticle formulations are known to be rapidly sequestered by the reticuloendothelial system (RES), primarily the liver, which commonly captures over 95% of injected nanoparticles (34, 35).

Our findings have important translational implications. While single-agent irinotecan is rarely curative for metastatic Ewing sarcoma (36), NV103 offers a platform for enhanced delivery of this otherwise potent agent. Moreover, the nanoparticle approach enables rational design of combinatorial therapies. Previous work has shown that doxorubicin-containing nanoparticles are also effective in Ewing models, although to a lesser extent than NV103. A combination of CD99-targeted nanoparticles carrying synergistic agents may yield further improvements in both the minimum effective dose (MED) and maximum tolerated dose (MTD) while minimizing systemic toxicity (37, 38).

This strategy opens up the possibility of reformulating frontline chemotherapy regimens such as VACIE (vincristine, doxorubicin, cyclophosphamide, ifosfamide, and etoposide) using targeted nanoparticles. Doing so could allow dose intensification without increased toxicity, ultimately improving survival outcomes while reducing long-term treatment-related morbidities. This is particularly relevant for pediatric and adolescent patients with Ewing sarcoma, who are at a high risk of late effects from conventional chemotherapy, including cardiotoxicity, infertility, and secondary malignancies.

In summary, the CD99-targeted NV103 platform demonstrated significant potential to improve both the efficacy and tolerability of chemotherapy in Ewing sarcoma. By achieving greater tumor kill at lower doses and with minimal toxicity, targeted nanoparticle delivery may represent a transformative approach to cancer treatment—one that addresses not only the imperative of cure but also the quality of life of survivors.

## Materials and Methods

### Composition of HPLNs

Hybrid Polymerized Liposomal Nanoparticles (HPLNs) were obtained from NanoValent Pharmaceuticals, Inc. (Bozeman, MT). The HPLNs were composed of N-(5’-hydroxy-3’-oxypentyl)-10-12-pentacosadiynamide (“h-Peg1-PCDA”), N-[methoxy(polyethylene glycol)-2000]-10-12-pentacosadiynamide (“m-Peg2000-PCDA”) (NanoValent Pharmaceuticals, Inc.), L-α-phosphatidylcholine hydrogenated soy (“hydrogenated soy PC”), 1,2-distearoyl-sn-glycero-3-phosphoethanolamine-N-[methoxy(polyethylene glycol)-2000] (“m-Peg2000-DSPE”), and cholesterol (Avanti Polar Lipids, Alabaster, Alabama). The antibody-conjugatable micelles were composed of m-Peg2000-DSPE and 1,2-distearoyl-sn-glycero-3-phosphoethanolamine-N-[maleimide(polyethylene glycol)-2000] (“mal-Peg2000-DSPE”).

### Preparation of HPLNs

Liposomes were formulated using hydrogenated soy PC, cholesterol, and m-Peg2000-DSPE in molar ratios of 57.5:37.5:5, respectively. HPLNs were prepared with hydrogenated soy PC, cholesterol, h-Peg1-PCDA, m-Peg2000-DSPE, and m-Peg2000-PCDA at molar ratios of 51:32:14:2:1. Antibody-conjugatable micelles were composed of m-Peg2000-DSPE and mal-Peg2000-DSPE in a 4:1 molar ratio. The preparation followed established protocols (29). Briefly, lipids were combined and evaporated under vacuum to form a thin film. This film was hydrated with 155 mM ammonium sulfate (pH 5.4) to achieve a 15 mM total lipid and cholesterol suspension. The suspension underwent bath sonication at 60°C for 60 minutes using a VWR Ultrasonic Cleaner Mod 250D. The resulting slightly opaque solution was filtered through 0.45 µm Supor Membrane syringe filter, 32 mm Acrodisc and extruded through stacked 80 nm polycarbonate membranes ten times using high pressure extruder (Avestin, Emulsiflex-C5 homogenizer) heated to 50°C. The nearly clear liposome solutions were cooled to 5°C and stored for 12 hours. After returning to room temperature, liposomes containing PCDA lipids were polymerized via UV irradiation at 254 nm using a Spectrolinker XL-1000 UV Crosslinker

(Spectronics Corp.) for 60 s with constant stirring. The ammonium sulfate solution exterior to the HPLNs was removed and buffer shifted to PBS (pH 7.4) on a Millipore Labscale TFF system with a Pellicon XL Cassette (Biomax 1000 kDA,). The resulting blue HPLNs were heated to 65°C for 5 minutes to convert them to the red (fluorescent) form. The solutions were then syringe-filtered through 0.2 µm Supor Membrane syringe filter, 32 mm Acrodisc to remove any insoluble contaminants.

### Irinotecan Loading

The HPLNs were incubated with irinotecan–HCl and heated to 60°C for 10 min. Unencapsulated Irinotecan was removed using a Millipore Labscale TFF system with a Pellicon XL Cassette (Biomax 1000kDA). The average particle size was measured using a Zetasizer Nano S (Malvern Instruments).

### Preparation of anti-CD99 antibody conjugated HPLNs

Human monoclonal IgG_1_ antibodies exclusively reactive with murine and human CD-99 (MIC2) antigen were prepared by Curia Global, Inc (Camararillo, California). Specificity and ability to bind to both fresh and formalin-fixed, paraffin embedded tissue were documented by both tissue immunohistochemistry of FFPE Ewing tumors from mice bearing human Ewing sarcomas and normal mouse tissues (heart, lung, liver, kidney, spleen), and FACS analysis of Ewing tumor cells (TC-32 and TC-71). The anti-CD99 antibodies were conjugated to the PEG termini of micelles using the maleimide-PEG coupling method (39). Briefly, thiolated (reduced) anti-CD99 antibodies were added to micelles composed of mal-Peg2000-DSPE and m-Peg2000-DSPE at a 1:4 molar ratio. The mixture was incubated at room temperature for 2 h to allow conjugation to maleimide residues on the micelles. Unbound maleimide residues were quenched with a 20 mM cysteine solution at room temperature for 30 min. Free cysteine and unbound antibodies were removed using Amicon Ultra Centrifugal Filters with a 100 kDa cutoff (Millipore). The anti-CD99 micelles were then incubated with the HPLNs at room temperature for 15 h. Following lipid insertion, the liposome–anti-CD99 micelle mixture was concentrated using Amicon Ultra 100 kDa centrifugal filters (Millipore) to obtain the final anti-CD99 micelle-HPLN conjugate (NV103).

### Quantification of entrapped liposomal Irinotecan

One hundred microliters of the sample was mixed with 880 µL of PBS buffer and 20 µL of 1% Triton X-100 solution in water to achieve a 1:10 dilution. The solution was then shaken at room temperature for 30 min at 850 RPM. Insoluble debris were removed from the sample by centrifugation at 24,000 × g for 5 min. The supernatant (500 µL) was loaded into a 3 kDa cutoff Centricon filter and spun at 5000 × g for ∼10 min. Twenty to 50 µl of the flow-through was transferred to a microplate and the volume was brought to 100 µL, noting the dilution factor. The absorbance was measured at 370 nm. The irinotecan concentration was calculated using a standard curve generated using free irinotecan.

### Tumor Cell Lines

TC-32 and TC-71 are Ewing sarcoma cell lines established by the corresponding author at the National Cancer Institute. Both possess the common EWSR::FLI1 fusion gene and express CD99. They have been maintained in the author’s lab and are part of the COG certified tumor cell line repository. Identity confirmation is based on periodic STR (short tandem repeat) assay and comparison with the parent low passage cell line splits maintained in the CHLA biorepository.

### Cell Viability Assay

The TC32 and TC71 Ewing sarcoma cell lines were cultured in RPMI 1640 medium supplemented with 10% fetal bovine serum (FBS) and antibiotics (penicillin/streptomycin). Cells were seeded into 96-well plates at a density of 1×10^4^ cells per well in 100 µL of medium and incubated overnight. The following day, cells were treated with anti-CD99-targeted irinotecan-loaded hybrid polymerized liposomal nanoparticles (NV103), untargeted irinotecan-loaded hybrid polymerized liposomal nanoparticles (HPLN/Ir), empty HPLNs, or free irinotecan for various durations, followed by washing with fresh media. Irinotecan concentrations ranged logarithmically from 0.01 to 100 μM, with a 0 nM control treated with phosphate-buffered saline PBS (pH 7.4). Each treatment was performed in triplicates. Cells were incubated under standard conditions (37°C, 5% CO₂) for 72 h. Cell viability was assessed using the CellTiter-Glo Luminescent Cell Viability Assay Kit (Promega, USA) according to the manufacturer’s protocol.

### Generation of Luciferase expressing cell lines

TC71 and TC32 (Ewing sarcoma cells) were transduced with the pCCL-MNDU3-LUC lentiviral vector encoding the firefly luciferase gene. Transduced cells were selected using 800 µg/mL G418. Luciferase activity was measured using the Luciferase Assay System (Promega), according to the manufacturer’s protocol.

### Injection of mice with luciferase-expressing tumor cells

TC71-Luc cells were cultured in RPMI 1640 medium supplemented with 10% fetal bovine serum (FBS) and antibiotics (penicillin/streptomycin). Prior to injection, cells were washed three times with phosphate-buffered saline (PBS), counted, and resuspended in serum-free antibiotic-free RPMI 1640 medium. Cell viability was assessed using a Vi-CELL XR Cell Viability Analyzer (Beckman Coulter), which employs the trypan blue dye exclusion method. Only preparations with over 90% viability were used for the injections. NOG (NOD.Cg-Prkdcscid Il2rgtm1Sug/JicTac) mice were obtained from Taconic and maintained under specific pathogen-free conditions at the Children’s Hospital, Los Angeles (CHLA) Animal Facility. All procedures adhered to the NIH Guidelines for Animal Care and were approved by the Institutional Animal Care and Use Committee (IACUC). For each 6–8-week-old female NOG mouse, 1-2 x10^6^ tumor cells were mixed with RPMI1640: Matrigel (1:1) and injected subcutaneously between the shoulder blades. Treatment began when the mean tumor size was a mean of 150 mm^3^All animal manipulations were performed under sterile conditions within a laminar flow hood.

### Bioluminescent imaging of the mice

Following cell injection, mice were imaged at various time points using an IVIS 100 bioluminescence/optical imaging system (Xenogen, Alameda, CA, USA). Ten minutes prior to imaging, D-luciferin (Xenogen) dissolved in PBS was administered intraperitoneally at a dose of 250 mg/kg. Anesthesia was induced with 5% isoflurane and maintained at 2.5% via a nose cone during the imaging. After capturing the photographic images, luminescent images were acquired with exposure times ranging from 1 to 60 s. The IVIS Living Image software (Xenogen) automatically superimposed grayscale and pseudocolor images to correlate the luciferase signals with their anatomical locations. Regions of interest (ROIs) were manually drawn around the bodies of the mice to assess the emitted signal intensity, expressed as photons per second within each ROI. Tumor bioluminescence correlated linearly with the tumor volume, as previously reported.

### Toxicity, immune response, and pathology studies

Female C57BL/6 mice, aged 6–8 weeks, were intravenously administered 10 mg/kg HPLN/Ir or NV103. Blood samples were collected 24 hours and two weeks post-injection through cardiac puncture using Microtainer tubes (Becton Dickinson) to separate the plasma. Whole blood was subjected to complete blood count (CBC) analysis, while plasma was utilized for clinical chemistry tests to assess liver and spleen function. The major organs were harvested, fixed in formalin, and processed for routine H&E staining. Imaging was performed using a Nikon epifluorescence microscope equipped with DP11 digital camera.

### Plasma Pharmacokinetic analyses

Irinotecan and its active metabolite, SN-38, were quantified in the plasma of C57BL/6 mice following a single intravenous administration of NV103 (5 mg/kg) and irinotecan (50 mg/kg). Blood samples were collected via cardiac puncture at specified time points under isoflurane anesthesia and immediately processed by centrifugation at 15,294 × g for 5 min at 4°C to isolate plasma.

Quantification was performed using a Waters Acquity UPLC I-Class system coupled with a Xevo TQS tandem quadrupole mass spectrometer utilizing TargetLynx v4.1 software. Chromatographic separation was achieved on a 2.1 × 50 mm, 1.7 µm BEH C18 column with a 0.4 mL/min flow rate. The mobile phase consisted of water and methanol, each containing 0.1% formic acid, in a gradient elution: 98% A / 2% B for 0.5 min, increasing to 98% B by 1.75 min, holding for 0.5 min, and returning to 98% A for 0.7 min.

Detection was performed in the positive electrospray ionization mode with multiple reaction monitoring (MRM). The transitions were m/z 587.2 → 124.1 for irinotecan, m/z 393.0 → 249.0 for SN-38, and m/z 455.3 → 150.1 for Verapamil (internal standard). Retention times were 1.63 min for irinotecan, 1.85 min for SN-38, and 1.74 min for verapamil, respectively.

Plasma calibration standards ranged from 2.5 to 1000 ng/mL and bone marrow calibrants were prepared by spiking blank samples. Protein precipitation was achieved using methanol with 0.1% formic acid followed by filtration through Ostro plates. Method validation adhered to the FDA guidelines to ensure linearity, accuracy, and precision.

Noncompartmental pharmacokinetic parameters were calculated, including area under the concentration-time curve (AUC₀–∞) using the trapezoidal rule, elimination half-life (t₁/₂ = 0.693/ke), total clearance (CLtot = dose/AUC), and apparent volume of distribution (Vd = dose / (AUC₀–∞ × ke)).

### Flow cytometry

To assess the binding efficiency of HPLNs, 1 × 10⁶ cells were incubated with 50 µg of HPLNs in 1 ml of culture medium for 1 h at room temperature. Subsequently, the cells were washed thrice with PBS, and the HPLN-bound cells were analyzed using a BD LSR II flow cytometer (BD Biosciences) equipped with a PE-Cy5 filter. Data acquisition was performed using the BD FACSDiva™ software (version 4.1.2). Unstained cells and cells treated with untargeted HPLNs were used as controls.

### Fluorescence Microscopy

To assess the binding and internalization of HPLNs in tumor cells, a 2-hour incubation at 37°C with 50 µg of HPLNs was performed. Following incubation, the cells were washed thrice with PBS and fixed with 2% formaldehyde at room temperature for 10 min. Mounting was done using VectaShield with DAPI (Vector Laboratories). Imaging was conducted using a Zeiss microscope equipped with a SPOT Insight QE Camera, employing a PE-Cy5 filter to visualize the bound nanoparticles.

### Protein lysates and Western blotting

The harvested cells were rinsed three times in PBS containing 100 mM Na_3_VO_4_ and then lysed in NP-40 Lysis buffer (50 mM HEPES, 100 mM NaF, 10 mM Na_4_P_2_O_7_, 2 mM Na_3_VO_4_, 2mM EDTA, 2 mM NaMoO_4_, and 0.5% NP-40) containing a commercial protease inhibitor cocktail (Roche) for 30 min at 4°C with shaking. After lysis/solubilization, protein concentrations were quantified using a DC Bio-Rad Protein assay kit (Bio-Rad Laboratories). After electrophoretic transfer to Immobilon-P membranes (Millipore), western blot analysis was performed using the indicated antibodies, including anti-CD99 antibodies (12EW) and anti-β-actin rabbit polyclonal antibody as the loading control.

### Support

This work was supported by an NCI grant, 1R44CA233128-01 Triche, Nagy (Co-PIs) 9/14/18-8/31/21, National Cancer Institute, NV103: Antibody Conjugated Nanoparticle for Ewing Sarcoma Targeted Therapy, NCI grant **1 R42 CA 281707-01,** Triche/Nagy, 08/2023 – 01/2026, Title: Phase II - Tumor Antigen Targeted Nanoparticle Therapy for Glioblastoma Multiforme (GBM), and Las Madrinas Endowment for Molecular Pathology.

### Disclaimers

Timothy Triche and Jon Nagy are co-founders of NanoValent and hold the founders’ stock in the company. None of the other authors declare any conflicts of interest.

## Author Contributions

Hyung-Gyoo Kang, Anahit Hovsepyran, Sheetal Mitra, and Jean Parmentier performed the animal experiments and the *in vitro* efficacy and imaging studies. Bryon Upton and Jon Nagy prepared the nanoparticles. Timothy Triche and Jon Nagy developed the nanoparticle technology. Timothy Triche conceived the project and wrote the manuscript with inputs from Hyung-Gyoo Kang, Sheetal Mitra, and Jean Parmentier.

## Acknowledgement

Pharmacokinetic and pharmacodynamic studies were performed by TD2 (Translational Drug Development, Scottsdale, Arizona).

Animal studies were performed with the support of the Small Animal Imaging Core, Saban Research Institute, Children’s Hospital, Los Angeles.

Cell imaging studies were performed with support from the Imaging Core, Saban Research Institute, Children’s Hospital, Los Angeles.

Nanoparticle characterization was performed in part by the Nanotechnology Core, Saban Research Institute, Children’s Hospital, Los Angeles.

## Data Access Statement

Research data supporting this publication are available from the corresponding author upon request.

## Supplemental Figures and Legends

**Supplement Figure 1.**
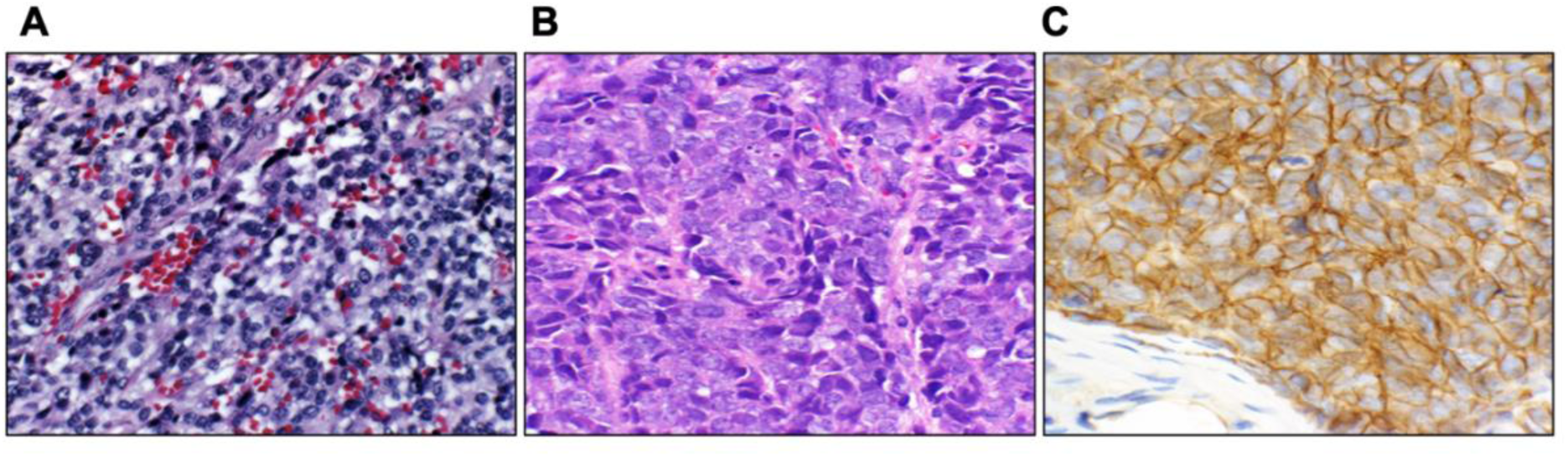
Establishment of a human Ewing Sarcoma xenograft mouse model. (A) H&E image of a subcutaneous flank tumor generated by implantation of human Ewing sarcoma cells into immunodeficient mice (B) H&E image of primary human Ewing (C) Immunohistochemical staining to show expression of CD99 on the surface of both xenograft and primary tumor cells

**Supplement Figure 2.**
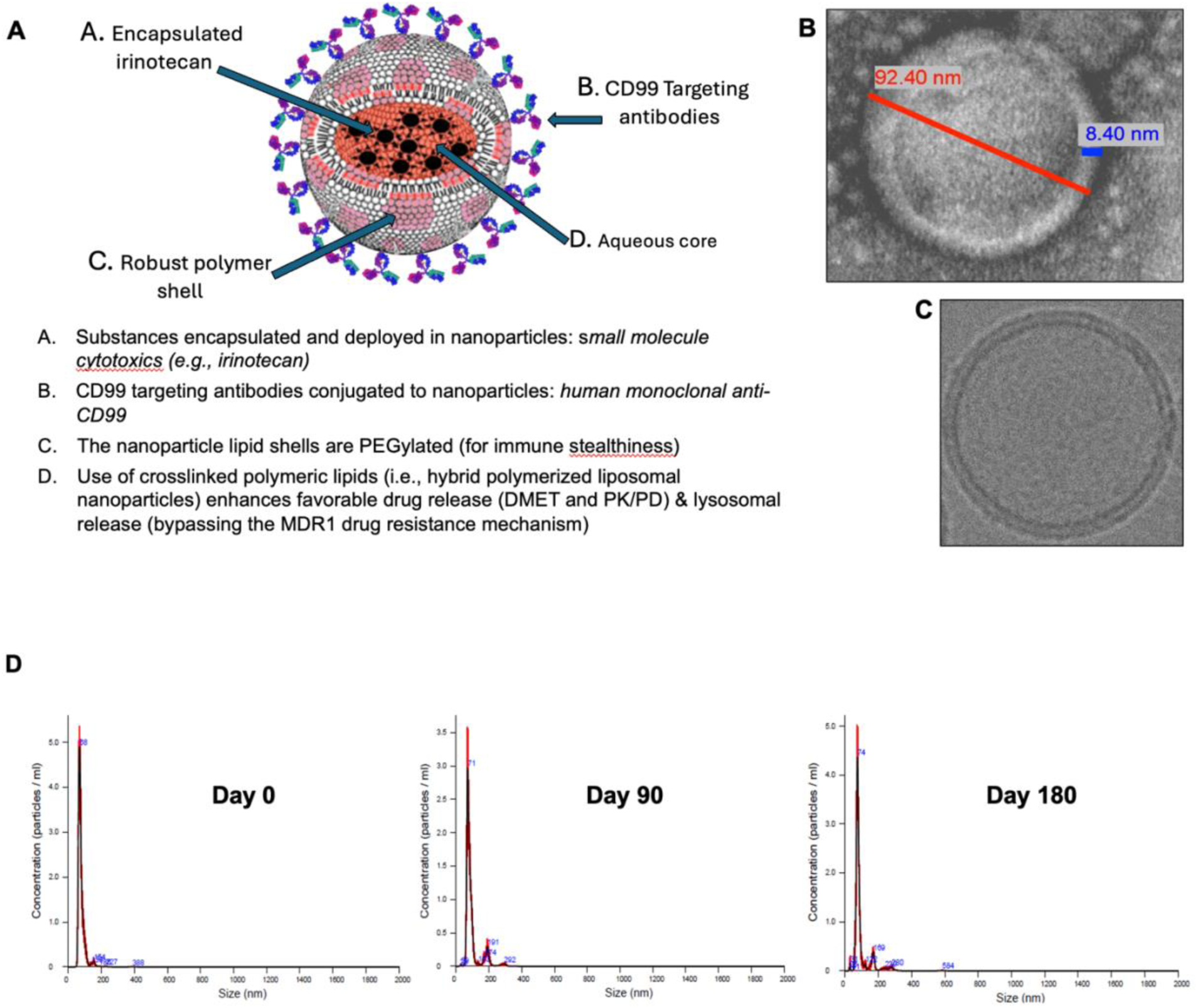
Schematic and Structural Characterization of Hybrid Polymerized Lipid Nanoparticles (HPLNs). (A) Diagram illustrating the composition of HPLNs, which consisted of cholesterol, hydroxy soy phosphatidylcholine (PC), pegylated distearoylphosphatidylethanolamine (DSPE), and pegylated 10,12-pentacosadiynoic acid (PCDA). Antibodies are conjugated to maleimide-terminated PEGylated DSPE micelles via reduced sulfhydryl bonds to form covalent linkages. (B) Transmission electron microscopy (TEM) image show the spherical morphology of nanoparticles, approximately 90 nm in diameter. (C) Cryo EM image shows the hollow aqueous core of nanoparticles.(D) Time-course NanoSight nanoparticle tracking analysis of NV103 size distribution.

**Supplement Figure 3.**
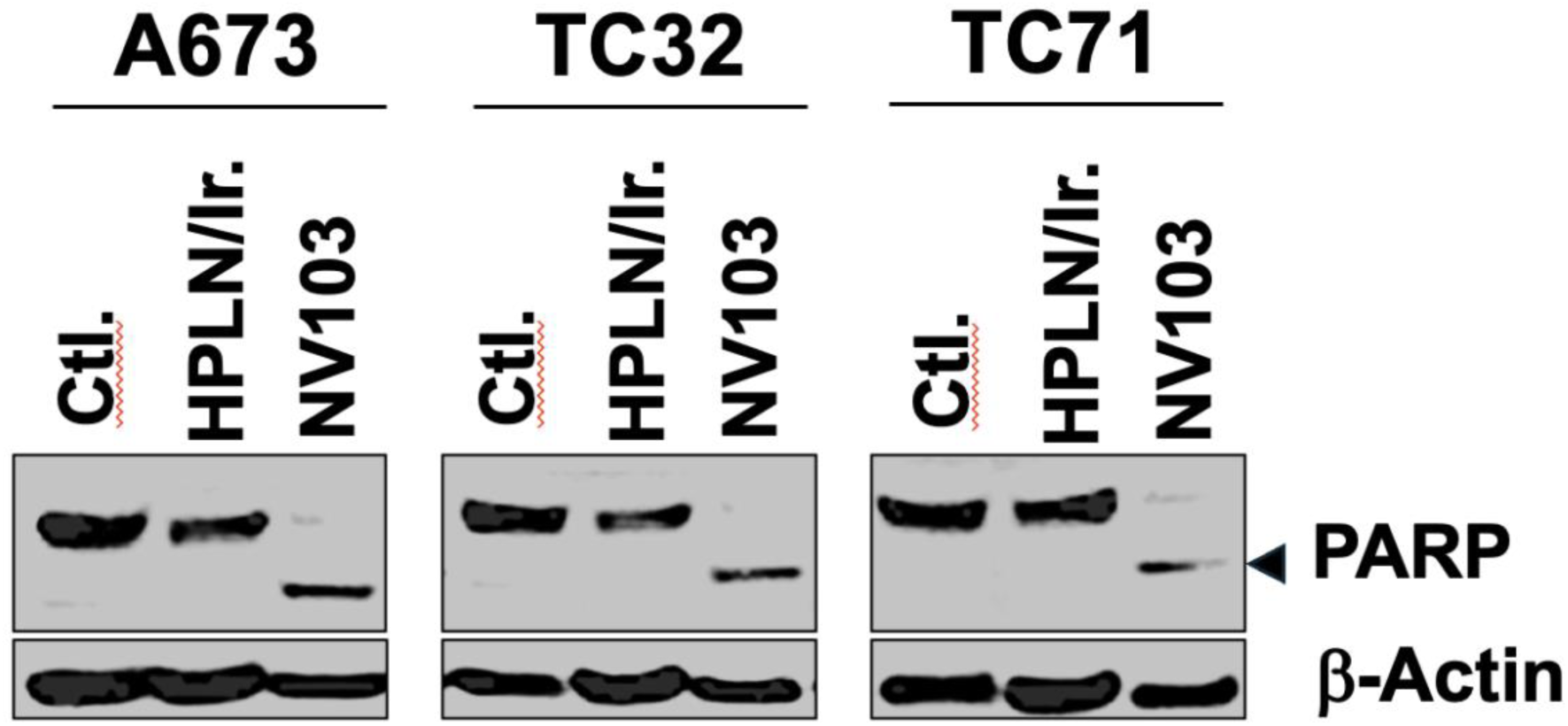
Induction of Apoptosis by NV103. A673 cells were treated with 5 mM HPLN/Ir or NV103 for two hours. The cells were then washed with fresh media, followed by 18 hours of culture. Lysates from treated cells were assayed for PARP cleavage by Western blotting using antibodies against PARP (arrowhead, cleaved PARP). β-Actin was used as a loading control.

**Supplement Figure 4.**
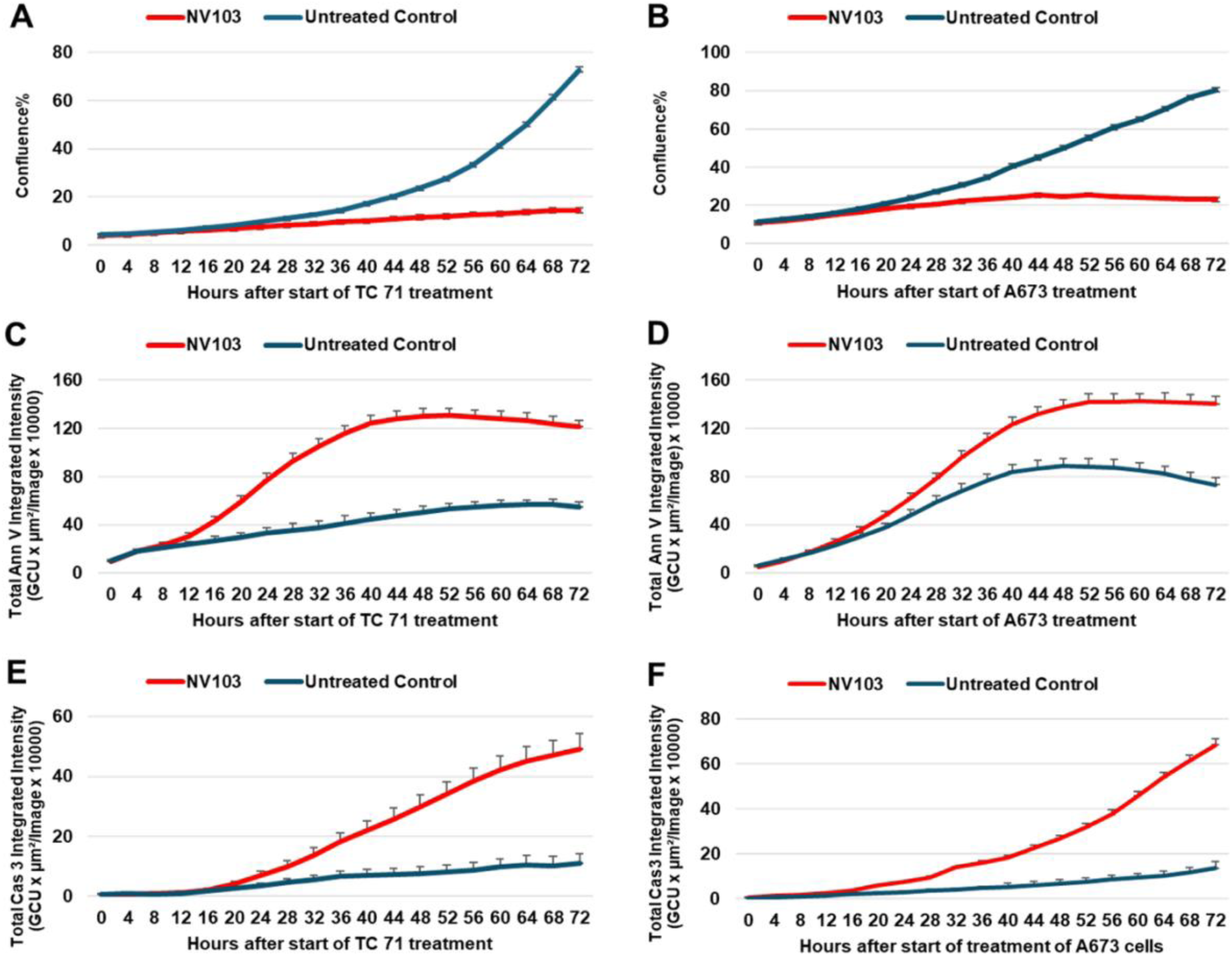
Onset of irinotecan-induced apoptosis in Ewing sarcoma cell lines following NV103 treatment using Incucyte. Cellular confluence percentage was measured over 72 h in TC71 (A) and A673 (B) Ewing sarcoma cells. Initiation of early apoptosis before 24 h as seen with detection of marker, Annexin V, in TC71 (C) and A673 (D) cells treated with nanoparticles. Late apoptosis was detected after 24 h using the caspase 3 assay in TC71 (E) and A673 (F) cells.

**Supplement Table 1.** Individual levels of Irinotecan and SN-38 in plasma following a single intravenous (IV) dose of NV103 at 5 mg/kg in C57BL/6 mice. BLQ: below limit of quantitation (LOQ=2.5 ng/mL) Samples were diluted as needed to fall within standard curve

| Dose mg/kg<br>Route | Animal<br>ID | Time<br>(hr) | Plasma<br>concentration<br>(ng/mL)<sup>a</sup> |  |
| --- | --- | --- | --- | --- |
|  |  |  | Irinotecan | SN-38 |
| Group 1<br>5 mg/kg<br>IV | 22 | 0.00 | BLQ | BLQ |
|  | 23 | 0.00 | BLQ | BLQ |
|  | 24 | 0.00 | BLQ | BLQ |
|  | 1 | 0.25 | 30600.00 | 5350.00 |
|  | 2 | 0.25 | 51000.00 | 8750.00 |
|  | 3 | 0.25 | 43050.00 | 12300.00 |
|  | 4 | 0.50 | 63700.00 | 8113.00 |
|  | 5 | 0.50 | 61550.00 | 8504.00 |
|  | 6 | 0.50 | 156000.00 | 5163.00 |
|  | 7 | 1.00 | 46150.00 | 11114.00 |
|  | 8 | 1.00 | 60750.00 | 8740.00 |
|  | 9 | 1.00 | 42100.00 | 9432.00 |
|  | 10 | 2.00 | 34850.00 | 10743.00 |
|  | 11 | 2.00 | 37150.00 | 11888.00 |
|  | 12 | 2.00 | 35200.00 | 8910.00 |
|  | 13 | 4.00 | 29450.00 | 911.60 |
|  | 14 | 4.00 | 14210.00 | 1490.00 |
|  | 15 | 4.00 | 28760.00 | 951.00 |
|  | 16 | 8.00 | 5000.00 | 974.00 |
|  | 17 | 8.00 | 8180.00 | 2525.60 |
|  | 18 | 8.00 | 9520.00 | 1330.80 |
|  | 19 | 24.00 | 420.00 | 38.40 |
|  | 20 | 24.00 | 310.00 | 985.60 |
|  | 21 | 24.00 | 980.00 | 97.20 |

**Supplement Table 2.** Summary of plasma levels of Irinotecan and SN-38 following a single IV dose of 5 mg/kg NV-103 (Group 1) in C57BL/6 mice. BLQ: below limit of quantitation (LOQ=2.5 ng/mL) NA: value not available

| Time (hr) | Irinotecan concentration (ng/mL) |  | SN-38 concentration (ng/mL) |  |
| --- | --- | --- | --- | --- |
|  | Ave | StdDev | Ave | StdDev |
| 0.00 | BLQ | NA | BLQ | NA |
| 0.25 | 41550.00 | 10282.39 | 8800.00 | 3475.27 |
| 0.50 | 93750.00 | 53920.80 | 7260.00 | 1826.55 |
| 1.00 | 49666.67 | 9809.73 | 9762.00 | 1220.92 |
| 2.00 | 35733.33 | 1239.29 | 10513.67 | 1502.19 |
| 4.00 | 24140.00 | 8606.55 | 1117.53 | 323.17 |
| 8.00 | 7566.67 | 2321.58 | 1610.13 | 812.64 |
| 24.00 | 570.00 | 359.30 | 373.73 | 530.71 |

**Supplement Table 3.** Individual levels of Irinotecan and SN-38 in plasma following a single IV dose of irinotecan, a standard agent, at 50 mg/kg in C57BL/6 mice. BLQ: below limit of quantitation (LOQ=2.5 ng/mL) Samples were diluted as needed to fall within standard curve

| Dose mg/kg<br>Route | Animal<br>ID | Time<br>(hr) | Plasma<br>concentration<br>(ng/mL)<sup>a</sup> |  |
| --- | --- | --- | --- | --- |
|  |  |  | Irinotecan | SN-38 |
| Group 2<br>50 mg/kg<br>IV | 22 | 0.00 | BLQ | BLQ |
|  | 23 | 0.00 | BLQ | BLQ |
|  | 24 | 0.00 | BLQ | BLQ |
|  | 1 | 0.25 | 25210.00 | 3849.00 |
|  | 2 | 0.25 | 28375.00 | 4366.00 |
|  | 3 | 0.25 | 20140.00 | 5165.00 |
|  | 4 | 0.50 | 23835.00 | 3172.00 |
|  | 5 | 0.50 | 28715.00 | 3166.00 |
|  | 6 | 0.50 | 30485.00 | 4165.00 |
|  | 7 | 1.00 | 15605.00 | 1545.00 |
|  | 8 | 1.00 | 18675.00 | 2706.00 |
|  | 9 | 1.00 | 22405.00 | 2637.00 |
|  | 10 | 2.00 | 11145.00 | 2872.00 |
|  | 11 | 2.00 | 7745.00 | 1599.00 |
|  | 12 | 2.00 | 9080.00 | 1602.00 |
|  | 13 | 4.00 | 958.00 | 241.20 |
|  | 14 | 4.00 | 1233.40 | 315.20 |
|  | 15 | 4.00 | 567.80 | 288.80 |
|  | 16 | 8.00 | 23.20 | 50.00 |
|  | 17 | 8.00 | 36.40 | 50.20 |
|  | 18 | 8.00 | 19.20 | 23.40 |
|  | 19 | 24.00 | 8.20 | 9.60 |
|  | 20 | 24.00 | BLQ | BLQ |
|  | 21 | 24.00 | BLQ | BLQ |

**Supplement Table 4.** Summary of plasma levels of Irinotecan and SN-38 following a single intravenous dose of irinotecan, a standard agent (Group 2), at 50 mg/kg in C57BL/6 mice. BLQ: below limit of quantitation (LOQ=2.5 ng/mL) NA: value not available

| Time (hr) | Irinotecan concentration (ng/mL) |  | SN-38 concentration (ng/mL) |  |
| --- | --- | --- | --- | --- |
|  | Ave | StdDev | Ave | StdDev |
| 0.00 | BLQ | NA | BLQ | NA |
| 0.25 | 24575.00 | 4154.06 | 4460.00 | 663.02 |
| 0.50 | 27678.33 | 3444.07 | 3501.00 | 575.05 |
| 1.00 | 18895.00 | 3405.33 | 2296.00 | 651.30 |
| 2.00 | 9323.33 | 1713.01 | 2024.33 | 734.10 |
| 4.00 | 919.73 | 334.45 | 281.73 | 37.50 |
| 8.00 | 26.27 | 9.00 | 41.20 | 15.42 |
| 24.00 | 8.20 | NA | 9.60 | NA |

